# Conservation of epigenetic regulation by the MLL3/4 tumour suppressor in planarian pluripotent stem cells

**DOI:** 10.1101/126540

**Authors:** Yuliana Mihaylova, Prasad Abnave, Damian Kao, Samantha Hughes, Alvina Lai, Farah Jaber-Hijazi, Nobuyoshi Kosaka, A. Aziz Aboobaker

## Abstract

Currently, little is known about the evolution of epigenetic regulation in animal stem cells. Using the planarian stem cell system to investigate the role of the COMPASS family of MLL3/4 histone methyltransferases, we demonstrate that their role as tumour suppressors in stem cells is conserved over a large evolutionary distance in animals. This also suggested the potential conservation of a genome wide epigenetic regulation program in animal stem cells, so we assessed the regulatory effects of Mll3/4 loss of function by performing RNA-seq and ChIP-seq on the G2/M planarian stem cell population, part of which contributes to the formation of outgrowths. We find many oncogenes and tumour suppressors among the affected genes that are therefore likely candidates for mediating MLL3/4 tumour suppression function in mammals, where little is known about *in vivo* regulatory targets. Our work demonstrates conservation of an important epigenetic regulatory program in animals and highlights the utility of the planarian model system for studying epigenetic regulation.

## Introduction

The pluripotent adult stem cell population of planarian flatworms is a highly accessible study system to elucidate fundamental aspects of stem cell function^1,2^. These stem cells, collectively known as neoblasts (NBs), bestow these animals with an endless capacity to regenerate all organs and tissues after amputation. Comparisons of stem cell expression profiles and functional data between animals show that some key aspects of stem cell biology are deeply conserved^3-9^, while others, like the transcription factors that define pluripotency in mammalian stem cells, appear not to be. Thus, studies of NBs have the potential to inform us about the origins of fundamental stem cell properties that underpin metazoan evolution, such as maintenance of genome stability^10^, self-renewal^7,11^, pluripotency^12-15^, differentiation^16-18^ and migration^19,20^. All of these are highly relevant to understanding human disease processes, particularly those leading to cancer.

Currently, very little comparative data exists for the role of epigenetic regulation in animal stem cells. Planarian NBs offer an opportunity to ask whether the cellular and physiological roles of different epigenetic regulators might be conserved between mammalian and other animal stem cells. Additionally, as mutations in many chromatin modifying enzymes are implicated in cancer^21-26^, using NBs as a model system may provide fundamental insight into why these mutations lead to cancer if epigenetic regulatory programs are conserved.

The genome-wide effects of chromatin modifying enzymes make understanding how they contribute to cancer phenotypes very challenging. Complexity in the form of tissue and cell heterogeneity, life history stage and stage of pathology make resolution of epigenetic regulatory cause and effect relationships *in vivo* very challenging. From this perspective, planarians and their easily accessible NB population may be a very useful model system. The planarian system could be particularly suitable for investigating the early transformative changes in stem cells at the onset of hyperplasia, as the NB identity of all potentially hyperplastic cells is known *a priori.* Here, we have employed this approach to study the planarian orthologs of the human tumour suppressors Mixed Lineage Leukaemia 3 (MLL3) and MLL4.

The human MLL proteins are the core members of the highly conserved COMPASS-like (<u>com</u>plex of <u>p</u>roteins <u>as</u>sociated with <u>S</u>et1) H3K4 methylase complexes. An extensive research effort has now established the evolutionary history and histone modifying activities of this protein family (**Supplementary Figure 1**^27-42^). Perturbation of MLL-mediated H3K4 methylase activity is characteristic of numerous cancer types. While prominent examples include the translocation events widely reported in leukaemias involving the *Mll1* gene^43-46^, the mutation rate of *Mll3* across malignancies of different origin approaches 7%, making *Mll3* one of the most commonly mutated genes in cancer^24^. In attempts to model the role of *Mll3* in cancer, mice homozygous for a targeted deletion of the *Mll3* SET domain were found to succumb to ureter epithelial tumours at high frequency^32^, an effect enhanced in a *p53*+/− mutational background. Heterozygous deletions of *Mll3* in mice also lead to acute myeloid leukaemia, as hematopoietic stem cells fail to differentiate correctly and over-proliferate, implicating *Mll3* in dose-dependent tumour suppression^26^. Recent studies have revealed an increasingly complicated molecular function of MLL3, its closely related paralog MLL4, and their partial *Drosophila* orthologs – *LPT* (<u>L</u>ost <u>P</u>HD-fingers of <u>t</u>rithorax-related; corresponding to the N-terminus of MLL3/4) and Trr (<u>t</u>rithorax-related; corresponding to the C-terminus of MLL3/4)^35,39^. LPT-Trr/MLL3/4 proteins have a role in transcriptional control via mono-methylating and/or tri-methylating H3K4 at promoters and enhancers^29,31,33-35,40,47,48^ (**Supplementary Figure 1**).

Links between mutations in *Mll3/4*, effects on downstream targets of MLL3/4 and human cancers remain to be elucidated. If the role of MLL3/4 in regulating stem cells is conserved in NBs, planarians could provide an informative *in vivo* system for identifying potential candidate target genes that drive tumour formation. Here we identify and investigate the role of *Mll3/4* orthologs in the planarian *Schmidtea mediterranea*, and show knockdown leads to NB over proliferation, tissue outgrowths containing proliferative NBs and differentiation defects. Investigating the regulatory effects accompanying this phenotype, we demonstrate mis-regulation of both oncogenes and tumour suppressors, providing a potential explanation for how tumour suppressor function is mediated by MLL3/4.

## Results

### The planarian orthologs of *Mll3/4* are expressed in stem cells

We found 3 partial orthologs of mammalian *Mll3* and *Mll4* genes. We named the planarian gene homologous to *Drosophila* LPT and the N-terminus of mammalian *Mll3/4* - *Smed-LPT* (KX681482) (**Supplementary Figure 2a**). Smed-LPT (LPT) protein contains two PHD-fingers and a PHD-like zinc-binding domain, suggesting that it has chromatin-binding properties^49^ (**Figure 1a**). There are two planarian genes homologous to *Drosophila* Trr and the C-terminus of mammalian *Mll3/4* – *Smed-trr-1* (KC262345) and *Smed-trr-2* (DN309269, HO004937), both previously described^36^. Both SMED-TRR-1 and SMED-TRR-2 contain a PHD-like zinc-binding domain, a FYRN (FY-rich N-terminal domain), FYRC (FY-rich C-terminal domain) and a catalytic SET domain. SMED-TRR-1 (TRR-1) contains only a single NR (Nuclear Receptor) box at a non-conserved position and SMED-TRR-2 (TRR-2) has no NR boxes (**Figure 1a**). This could indicate some functional divergence exists between TRR-1 and TRR-2, where only TRR-1 is capable of interacting with nuclear receptors.^33,35,50,51^.

**Figure 1.**
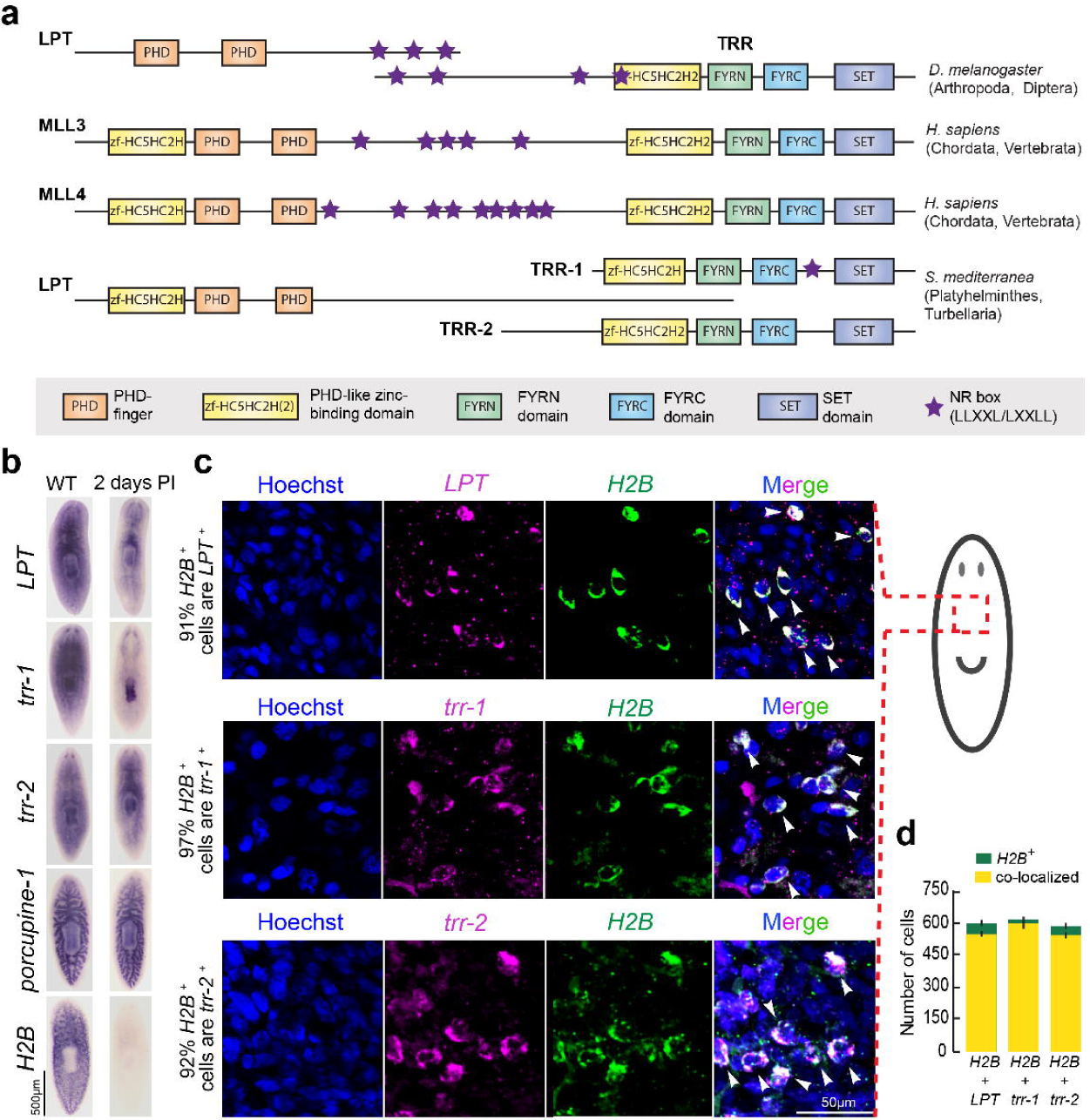
S. *mediterranea* has three partial *Mll3/4* orthologs expressed in stem cells. **(a)** A schematic depicting the structure and domain composition of MLL3/MLL4 proteins in *D. melanogaster, H. sapiens* and *S. mediterranea.* **(b)** genes’ expression pattern in wildtype (WT) and two days following a lethal dose (60 Gy) of gamma irradiation (PI = post-irradiation). *Porcupine-1* (expressed in the irradiation-insensitive cells of the differentiated gut) and *H2B* (expressed in the irradiation-sensitive neoblasts) are used as a negative and positive control respectively. Ten worms per condition were used. **(c)** White arrows point to examples of cells double-positive for *Mll3/4* transcripts and *H2B* transcripts. The schematic shows the body area imaged. **(d)** Graph showing the raw cell counts used for percentage estimates in (c). Green colour represents all counted *H2B*-positive cells, yellow represents H2B-positive cells also expressing an *Mll3/4* ortholog. Error bars represent Standard Error of the Mean (SEM). Ten animals per condition were used.

We performed whole-mount *in situ* hybridization (WISH) and found that *LPT, trr-1* and *trr-2* are broadly expressed across many tissues and organs. Gamma irradiation to remove cycling cells in S. *mediterranea* revealed that the three transcripts are also likely to be expressed in NBs (**Figure 1b**). A double fluorescent *in situ* hybridization (FISH) with the pan-stem cell marker *Histone 2B (H2B)* confirmed that over 90% of all NBs co-express *LPT, trr-1* and *trr-2* (**Figure 1c-d**). The genes also showed clear expression in the brain, pharynx and other differentiated tissues (**Figure 1b**).

### Loss of *Mll3/4* function leads to regeneration defects and tissue outgrowths

In order to study the function of planarian *Mll3/4*, we investigated phenotypes after RNAi-mediated knockdown. Following LPT(RNAi), there was a clear failure to regenerate missing structures, including the eyes and pharynx, with regenerative blastemas smaller than controls (**Figure 2a-b**). After 7 days of regeneration we observed that, as well as failure to regenerate missing structures, animals began to form tissue outgrowths (**Figure 2c**). Intact (homeostatic) *LPT(RNAi)* animals also developed outgrowths, but at a lower frequency than regenerates (**Figure 2d**).

**Figure 2.**
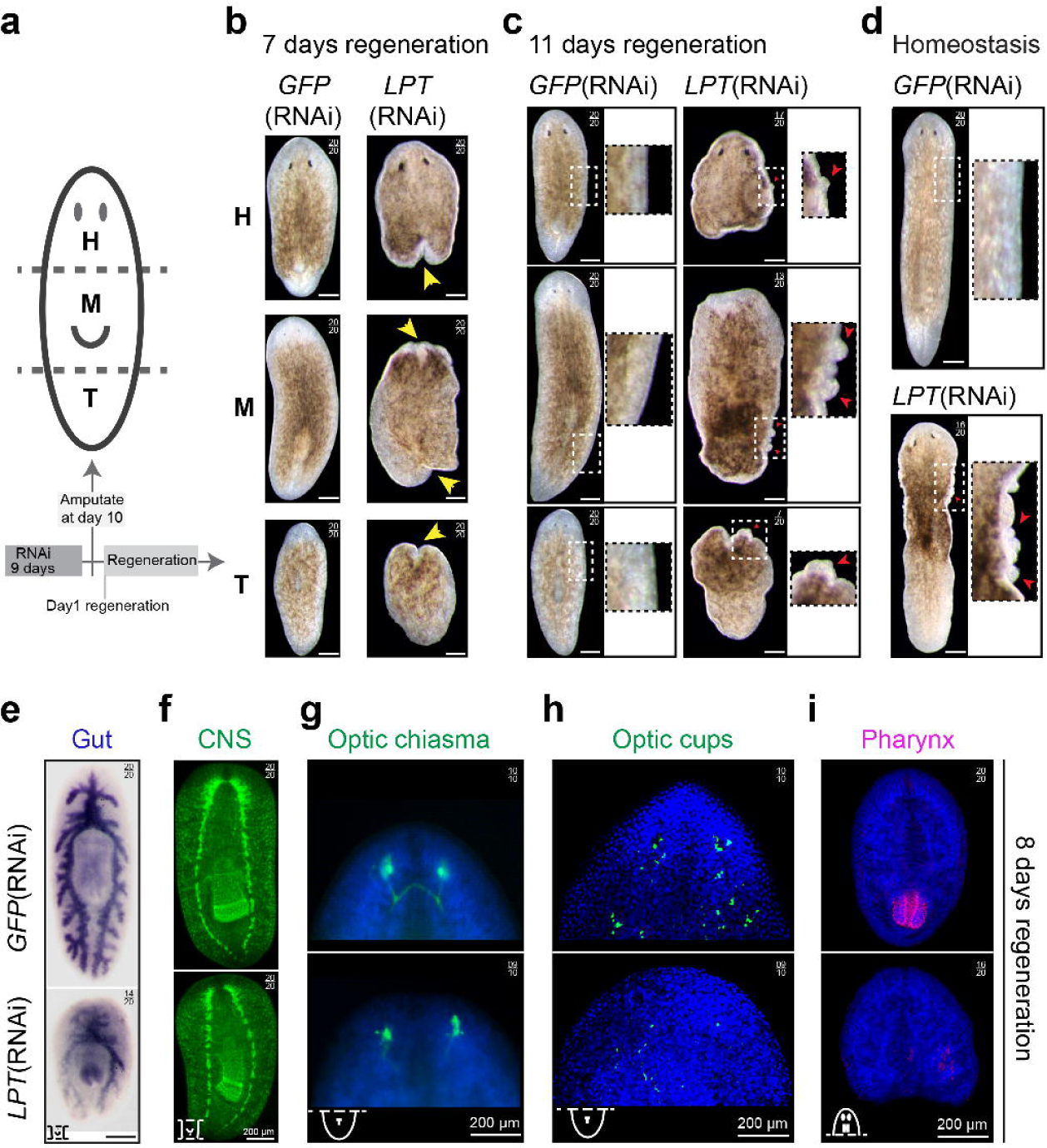
LPT(RNAi) results in differentiation defects and outgrowth formation during regeneration. **(a)** A schematic showing the amputation of RNAi worms into head (H), middle (M) and tail (T) pieces in order to observe regeneration of different structures. The time-course of all the experiments on *Mll3/4* knockdown animals is depicted underneath the worm schematic. A total of 9 days of dsRNA microinjection-mediated RNAi was followed by amputation on the 10^th^ day and subsequent observation of regeneration. **(b)** Head, middle and tail pieces following *LPT*(RNAi) or control GFP(RNAi) at day 7 of regeneration. Yellow arrows point towards the defects in blastema formation. **(c)** Head, middle and tail pieces following *LPT*(RNAi) or control GFP(RNAi) at day 11 of regeneration. Red arrows point towards outgrowths. **(d)** Homeostatic animals following *LPT*(RNAi) or control GFP(RNAi) at day 14 post RNAi. Red arrows point towards outgrowths. **(e)** Gut regeneration and maintenance in middle pieces following *LPT*(RNAi), as illustrated by RNA probe for the gene *porcupine-1* at 8 days of regeneration. **(f)** Brain regeneration in middle pieces at 8 days post-amputation following *LPT*(RNAi), as illustrated by anti-SYNORF-1 antibody labeling the central nervous system (CNS). **(g)** Optic chiasm recovery in tail pieces at 8 days of regeneration following *LPT*(RNAi), as shown by anti-VC-1 antibody. **(h)** Recovery of optic cups and organized trail of optic cup precursor cells in tail pieces at 8 days of regeneration following *LPT*(RNAi), as demonstrated by RNA probe for *SP6-9.* **(i)** Pharynx recovery in head pieces at 8 days of regeneration following *LPT*(RNAi), as illustrated by RNA probe for *laminin.*

Following individual knockdown of *trr-1* and *trr-2*, milder differentiation defects were observed compared to *LPT*(RNAi), with no obvious outgrowths (**Supplementary Figure 2b-d**), confirming results from an earlier study^36^. However, *trr-1/trr-2* double knockdown recapitulated the phenotype of *LPT*(RNAi), but with higher penetrance and increased severity (**Supplementary Figure 2e,f**). Functional redundancy between the two *trr* paralogs likely accounts for the reduced severity after individual knockdown. Double knockdown animals all developed outgrowths and started dying at day 5 post-amputation. Based on these observations, we decided to focus our attention on the *LPT*(RNAi) phenotype as regeneration defects and the formation of tissue outgrowths were temporally distinct and could be studied consecutively.

A more thorough study of the differentiation properties of *LPT*(RNAi) animals following amputation showed that the triclad gut structure failed to regenerate secondary and tertiary branches and to extend major anterior and posterior branches (**Figure 2e**). Cephalic ganglia (CG) regenerated as smaller structures, the two CG lobes did not join in their anterior ends in *LPT*(RNAi) animals (**Figure 2f**) and the optic chiasm and optic cups were mis-patterned and markedly reduced (**Figure 2g-h**). We found that 80% of *LPT*(RNAi) animals did not regenerate any new pharyngeal tissue (**Figure 2i**). We interpreted these regenerative defects as being indicative of either a broad failure in stem cell maintenance and/or differentiation.

### LPT/trr-1/trr-2 function is required for correct differentiation of epidermal and neural lineages

One of the structures most severely affected following loss of *LPT* function was the brain so we looked at the regeneration of different neuronal subtypes. **LPT*(RNAi)* animals had reduced numbers of GABAergic (**Figure 3a**), dopaminergic (**Figure 3b**), acetylcholinergic (**Figure 3c**) and serotonergic (**Figure 3d**) neurons. Brain defects were milder following knockdown of either *trr-1* or *trr-2* in agreement with the hypothesis of some functional redundancy between these paralogs (**Supplementary Figure 3a-d**).

**Figure 3.**
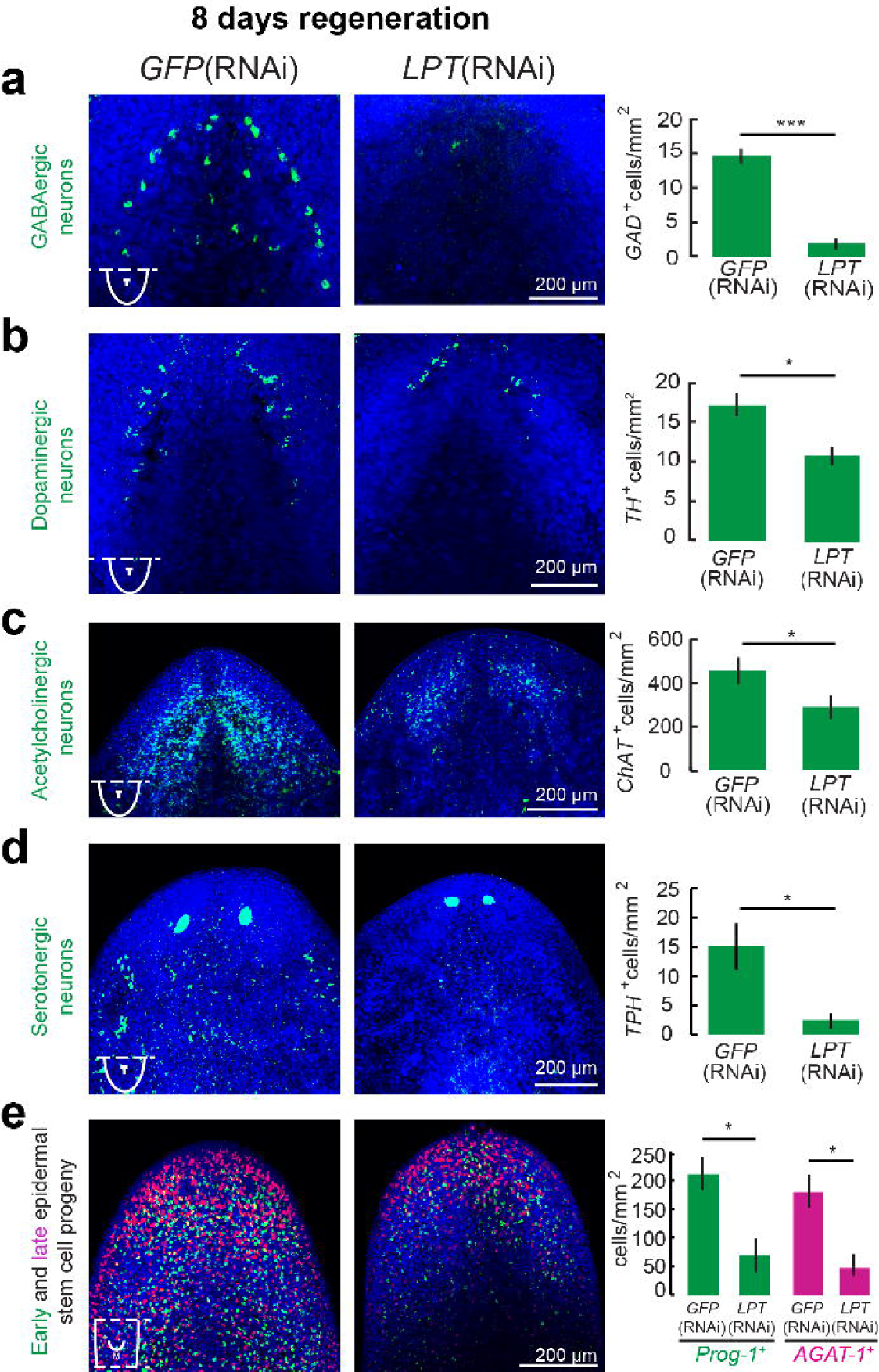
LPT controls differentiation across neuronal and epidermal lineages. **(a)** Quantification of the number of GABAergic neurons (labeled by *GAD)*, **(b)** dopaminergic neurons (labeled by *TH)*, **(c)** acetylcholinergic neurons (labeled by *ChAT)*, **(d)** serotonergic neurons (labeled by *TPH)* and **(e)** early (labeled by *prog-1*) and late (labeled by *AGAT-1*) epidermal stem cell progeny at 8 days of regeneration following *LPT*(RNAi). 2-tailed t-test was used for all comparisons; *p<0.05, ***p<0.001. Error bars represent Standard Error of the Mean (SEM). Ten animals per condition per experiment were assessed over the course of two separate experiments.

Epidermal tissue was also affected. Both early *(Prog-1*^*+v*^ cells) and late (AGAT-1^+ve^ cells) epidermal progeny were significantly decreased, but not entirely absent, in *LPT*(RNAi) regenerating animals (**Figure 3e**). No such defect was seen in individual *trr-1* and *trr-2* knockdown animals (**Supplementary Figure 3e**).

These effects along with defects in pharynx and gut tissues implicate broad NB differentiation defects in **LPT*(RNAi)* animals.

### LPT limits normal stem cell proliferation and tissue growth

Aside from impairment of regeneration following *LPT*(RNAi), the other major phenotype we observed were outgrowths of tissue that appeared at unpredictable positions in regenerating pieces.

Planarian regeneration is characterized by an early burst of increased NB proliferation, 6-12 hours after wounding, and a second peak of proliferation, 48 hours after amputation^52^. Following *LPT*(RNAi), we observed significant increases in proliferation at both of these peaks and at 8 days postamputation, as proliferation failed to return to normal homeostatic levels (**Figure 4a**). It was previously reported that *Trr-1*(RNAi) leads to decreases in mitotic cells, whereas *Trr-2*(RNAi) doesn’t affect NB proliferation^36^.

**Figure 4.**
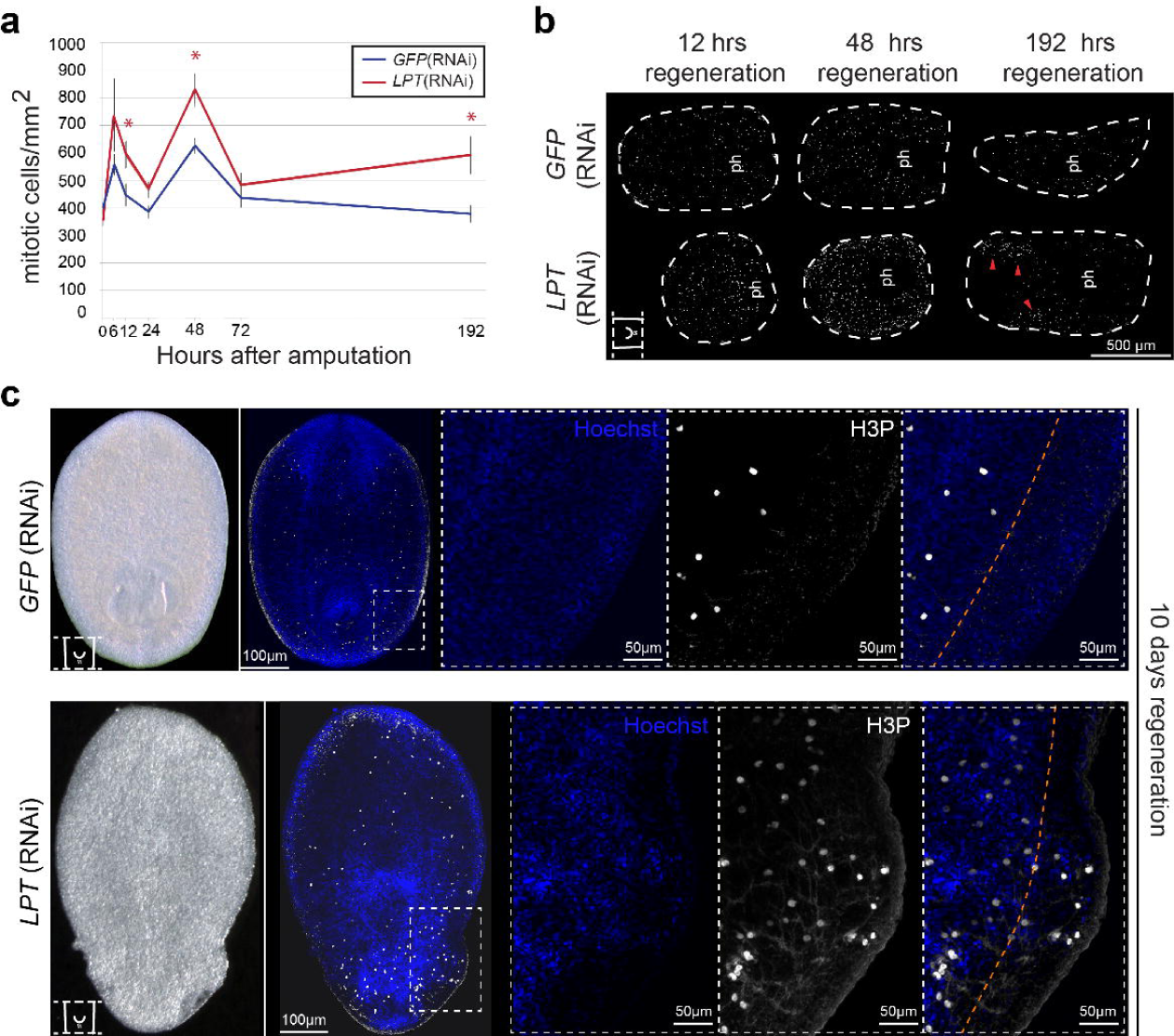
Over-proliferation and mitotic cell clustering precedes and accompanies the emergence of outgrowths in *LPT(RNAi*) regenerating animals. **(a)** Quantification of mitotic cells (labeled by anti-H3P antibody) at different post-amputation time-points following *LPT*(RNAi). N=10 animals per time-point. 2-tailed t-test was used for analysis; *p<0.05. Error bars represent Standard Error of the Mean (SEM). **(b)** Examples of middle pieces at the time-points post-amputation showing significant difference in mitotic cell (white) counts according to **(a)**. ‘ph’ indicates the pharynx. The red arrows point towards clusters of mitotic cells in late stage regenerates (192 hrs./8 days). **(c)** Mitotic cells (white) are always restricted to mesenchyme tissue in GFP(RNAi) animals but are penetrated into epidermal outgrowths in *LPT*(RNAi) animals. Orange line indicates the tentative border of mesenchyme.

In 8 day-regenerating *LPT*(RNAi) worms the observed over-proliferation is a result of localized clusters of mitotic cells rather than broad increase in proliferation across regenerating animals (**Figure 4b**). This is different from previously reported planarian outgrowth phenotypes from our group and others, where hyperplastic stem cells are evenly distributed^53,54^. It seems likely that these mitotic clusters might eventually be responsible for the formation of outgrowths. When we looked at outgrowths, we found mitotic cells usually restricted to mesenchymal tissue, had penetrated into epidermal outgrowths in *LPT*(RNAi) animals (**Figure 4c**).

In order to understand if ectopically cycling NBs represented the breadth of known stem cell heterogeneity in planarians or only a subset of lineages, we performed FISH for markers of the *sigma* (collectively pluripotent NBs), *zeta* (NBs committed to the epidermal lineage) and *gamma* (NBs committed to the gut lineage) cell populations^55^. We found that all three NB populations are represented in the outgrowths of *LPT*(RNAi) animals (**Figure 5a-c**). *Sigma, zeta* and *gamma* NBs are not significantly increased in pre-outgrowth *LPT*(RNAi) animals (**Supplementary Figure 4**), suggesting that the presence of these cells in outgrowths is not a secondary effect of increased cell number and passive spread of these cell populations, but rather local proliferation at outgrowth sites.

**Figure 5.**
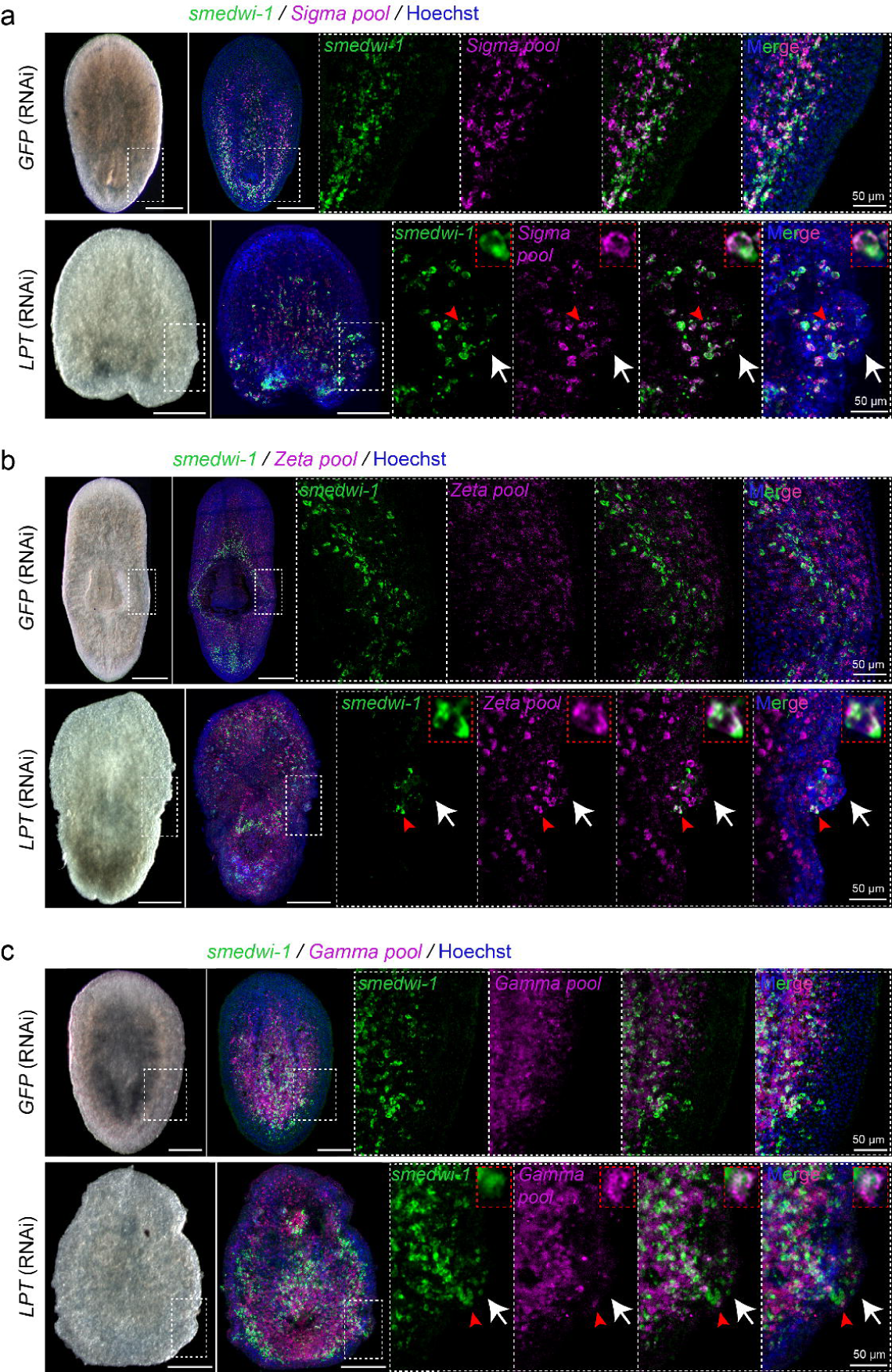
Pluripotent as well as lineage restricted stem cells are present in outgrowths of *LPT*(RNAi) animals. **(a)** Head piece showing distribution of *Sigma* stem cells in GFP(RNAi) and *LPT*(RNAi) animals. *Sigma* stem cells are double positive for *smedwi-1* and the *‘Sigma pool’* of RNA probes *(Soxp1, Soxp2).* White arrows in *LPT*(RNAi) animals point towards the outgrowth. Red arrows indicate a double-positive cell magnified in red inset box. Scale bars: 200 µm. **(b)** Images showing distribution of *Zeta* stem cells in GFP(RNAi) and *LPT*(RNAi) animals. *Zeta* stem cells are double positive for *smedwi-1* and the *‘Zeta pool’* of RNA probes *(zfp-1, Soxp3, egr-1).* White arrows in *LPT*(RNAi) animals point towards the outgrowth. Red arrows indicate a double-positive cell magnified in red inset box. Scale bars: 200 µm. **(c)** Images showing distribution of *Gamma* stem cells in GFP(RNAi) and *LPT*(RNAi) animals. *Gamma* stem cells are double positive for *smedwi-1* and the *‘Gamma pool* of RNA probes *(gata4/5/6, hnf4).* White arrows in *LPT*(RNAi) animals point towards the outgrowth. Red arrows indicate a double-positive cell magnified in red inset box. Scale bars: 200 µm.

The epidermal progeny, marked by *Prog-1* and *AGAT-1*, were concentrated in the outgrowths of *LPT*(RNAi) animals, while being in reduced numbers in nonoutgrowth tissue (**Supplementary Figure 5a**). The observed disarray of *Prog-1*^*+*^ and *AGAT-1*^*+*^ cells in outgrowths could be the result of perturbed patterning and polarity of the epidermal layer in *LPT*(RNAi) animals (**Supplementary Figure 5b**), as epidermal cells appear to have lost polarity and to be no longer capable of forming a smooth epidermal layer. Furthermore, the average epidermal nuclear size is significantly increased in **LPT*(RNAi)* animals compared to controls (**Supplementary Figure 5c**), an effect similar to the pathology seen following knockdown of the tumour suppressor *SMG-1*^*54*^. The epithelial layer in *LPT*(RNAi) animals also appears less well-defined than that in control animals, with a blurred distinction between epithelium and mesenchyme. Another feature of the *LPT*(RNAi) phenotype, encountered in a variety of malignancies^56^, are changes in nuclear shape (**Supplementary Figure 5d**).

In summary, LPT controls NB proliferation and restricts stem cells to predefined tissue compartments as well as being responsible for the successful differentiation of several lineages. Taken together, our data demonstrate that disturbance of the function of planarian LPT leads to development of both differentiation and proliferation defects (**Figure 6**), allowing us to conclude that the function of LPT/trr/Mll3/4 proteins as an epigenetic tumour suppressor function is conserved over a large evolutionary distance.

**Figure 6.**
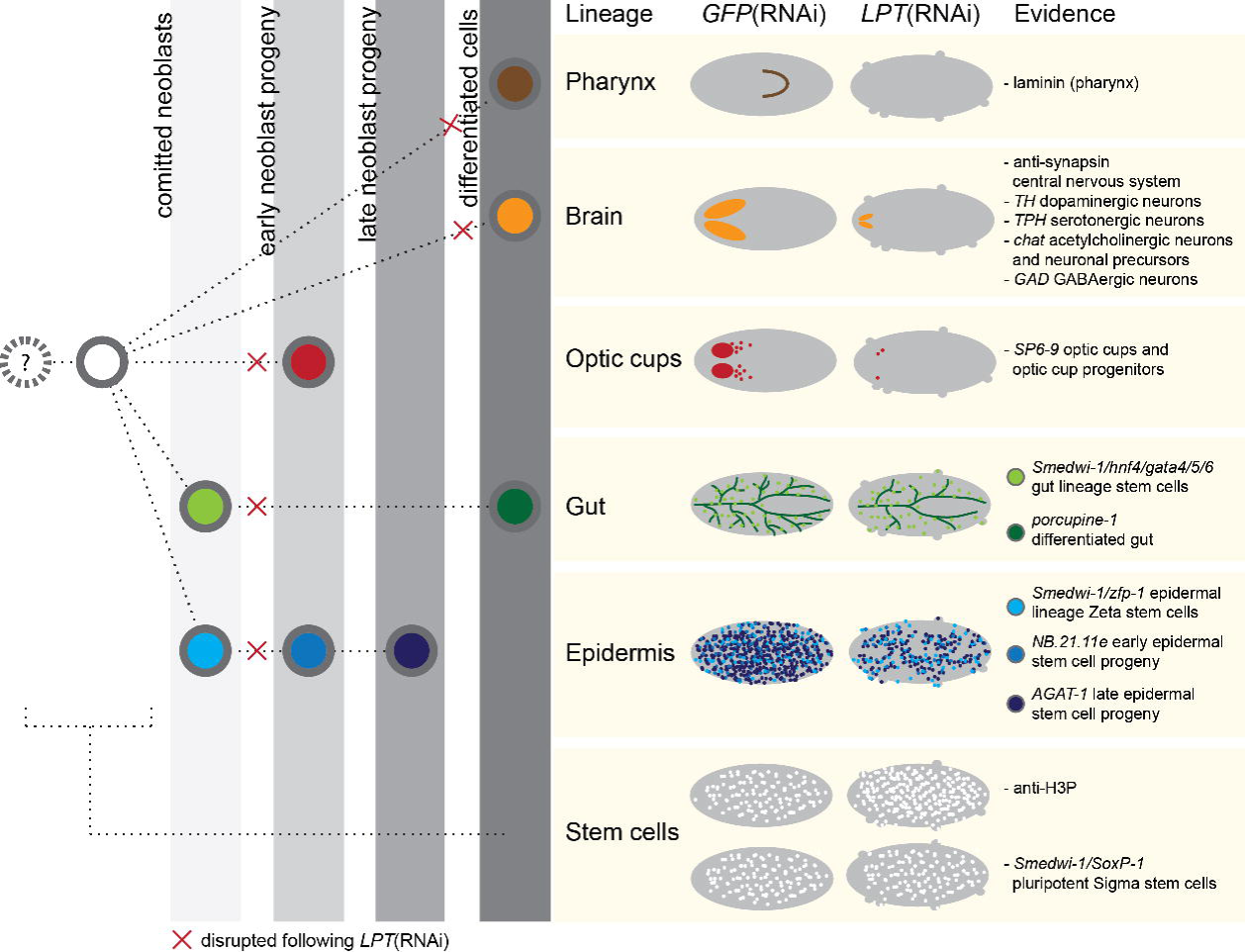
*LPT*(RNAi) results in a cancer-like phenotype. A summary of the differentiation and neoblast proliferation data presented, together with a simplified flowchart illustrating the tested lineages’ development under knockdown conditions. A red cross sign indicates where the defect in a lineage is detected following *LPT*(RNAi).

### *LPT*(RNAi) results in transcriptional changes consistent with driving proliferation in stem cells

A key insight missing from the literature for *Mll3* and *Mll4* mutations, is the downstream targets that are mis-regulated in disease states, for example in hematopoietic stem cells that cause leukaemia^26^. Given the conserved tumour suppressor function in planarians, we decided to focus on early regeneration when *LPT*(RNAi) animals that do not yet exhibit any outgrowth phenotype, providing the possibility to describe early regulatory changes that are potentially causal of out growths, rather than consequential. We performed RNA-seq on X1 (G2/M) fluorescence activated cell sorted (FACS) NBs from *LPT*(RNAi) and GFP(RNAi) planarians at 3 days of regeneration. Our analysis revealed that 540 transcripts are down-regulated (fold change <= −1.5, p<0.05) and 542 -up-regulated (fold change >= 1.5, p<0.05) in X1 stem cells from *LPT*(RNAi) animals when compared to controls (**Supplementary Table 1**).

A recent meta-analysis of all available S. *mediterranea* RNA-seq data allowed classification of all expressed loci in the planarian genome by their relative expression in FACS sorted cell populations representing stem cells, stem cell progeny and differentiated cells^9^. Superimposing the differentially expressed genes following *LPT*(RNAi) onto a gene expression spectrum reflecting FACS compartments, shows that *LPT*(RNAi) has a broad effect on gene expression in X1 cells (**Figure 7a**), affecting genes normally expressed in different planarian cell types (**Figure 7b**).

**Figure 7.**
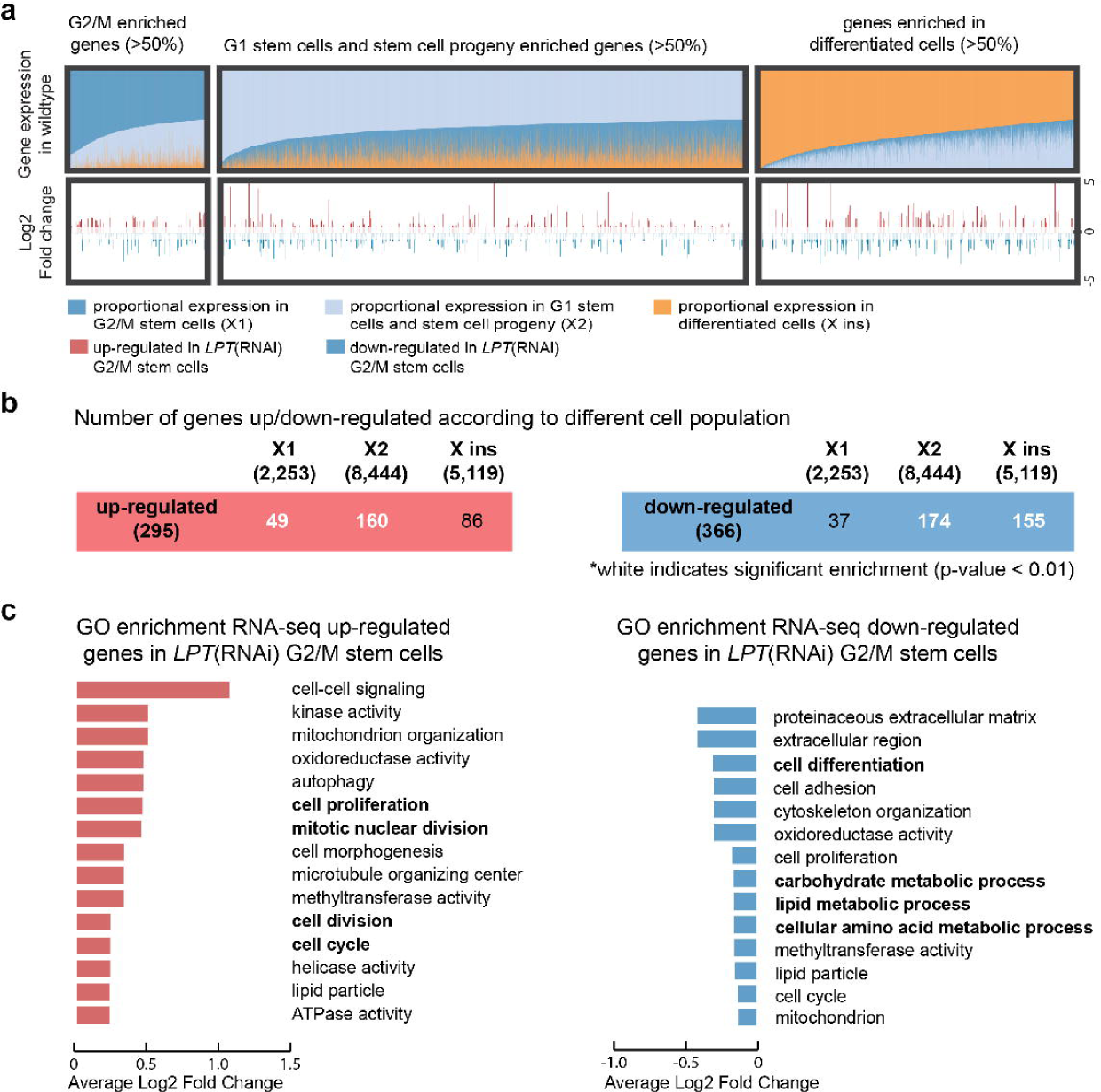
RNA-seq of G2/M stem cells following *LPT*(RNAi) reveals effects on genes enriched in different cell populations. **(a)** Genes were classified according to their proportional expression in the X1 (G2/M stem cells; dark blue), X2 (G1 stem cells and stem cell progeny; light blue) and X ins (differentiated cells; orange) FACS populations of cells. Genes were defined as enriched in certain population(s) if more that 50% of their expression is observed in that population in wild type animals. Each vertical line represents a gene. Under the population expression enrichment track is a track with all the significantly up- and down-regulated genes in G2/M stem cells following *LPT*(RNAi). The genes with fold change >1.5 (p<0.05) are shown in red following a log2 fold change transformation. The genes with fold change <-1.5 (p<0.05) are shown in blue following a log2 fold change transformation. The Wald’s test (as part of the Sleuth software) was used for assessing differential expression. **(b)** Enrichment for genes in each of the three classes was calculated for the up- and down-regulated genes’ list (red and blue respectively). The number of genes in each group is indicated in brackets under the group’s name. Numbers in white represent significant enrichment (p<0.01) according to a hypergeometric enrichment test. **(c)** Gene Ontology (GO) enrichment analysis on the genes significantly up-regulated (red) and down-regulated (blue) in G2/M stem cells following *LPT*(RNAi). Categories are sorted by average Log2 fold change of the up- or down-regulated genes falling in each category. In bold are shown terms that relate to the described *Mll3/4* loss of function phenotype.

Analysis of Gene Ontology (GO) terms revealed a clear enrichment for cell cycle and cell division-associated terms in the list of up-regulated genes (**Figure 7c**), in agreement with the observed hyper-proliferation in *LPT*(RNAi) phenotype. The list of down-regulated genes is also enriched for cell cycle-related terms, as well as cell differentiation and metabolism-related processes (**Figure 7c**). These findings suggest a broad link between gene expression changes caused by LPT loss of function and NB over-proliferation.

### *LPT*(RNAi)-induced changes to promoter H3K4 methylation and transcription are correlated

Previous studies tie MLL3/4/LPT-Trr function directly to mono- and tri-methylation of H3K4^29,31-34^ and indirectly to tri-methylation of H3K27, because the H3K27me3 demethylase UTX is present in the same protein complex^57^. We set out to understand potential epigenetic causes of the transcriptional changes following *LPT*(RNAi) in planarians. To this end, we performed Chromatin Immunoprecipitation sequencing (ChIP-seq) on isolated planarian G2/M stem cells and used Drosophila S2 cells as a spike-in control to normalize for any technical differences between samples^58^. The profile of H3K4me3, H3K4me1 and H3K27me3 at the transcriptional start sites (TSSs) of genes in control X1 cells was in agreement with their conserved roles in transcriptional control^9,34^ (**Figure 8a, Supplementary Figure 6**), suggesting out methodology was robust. *LPT*(RNAi) led to only subtle changes in the overall level of H3K4me3 and H3K4me1 at TSSs throughout the genome. A decrease in H3K4me1and H3K4me3 was apparent proximal to the TSS in genes where expression is normally enriched in differentiated cells (**Figure 8a**). Concomitant with this, we also observed an increase in H3K4me1 signal downstream of the predicted TSS for genes in enriched in stem cells (**Figure 8a**). For the H3K27me3 mark, no consistent pattern was observed as a result of *LPT*(RNAi) in any group of genes subdivided by expression profiles (**Figure 8a**).

**Figure 8.**
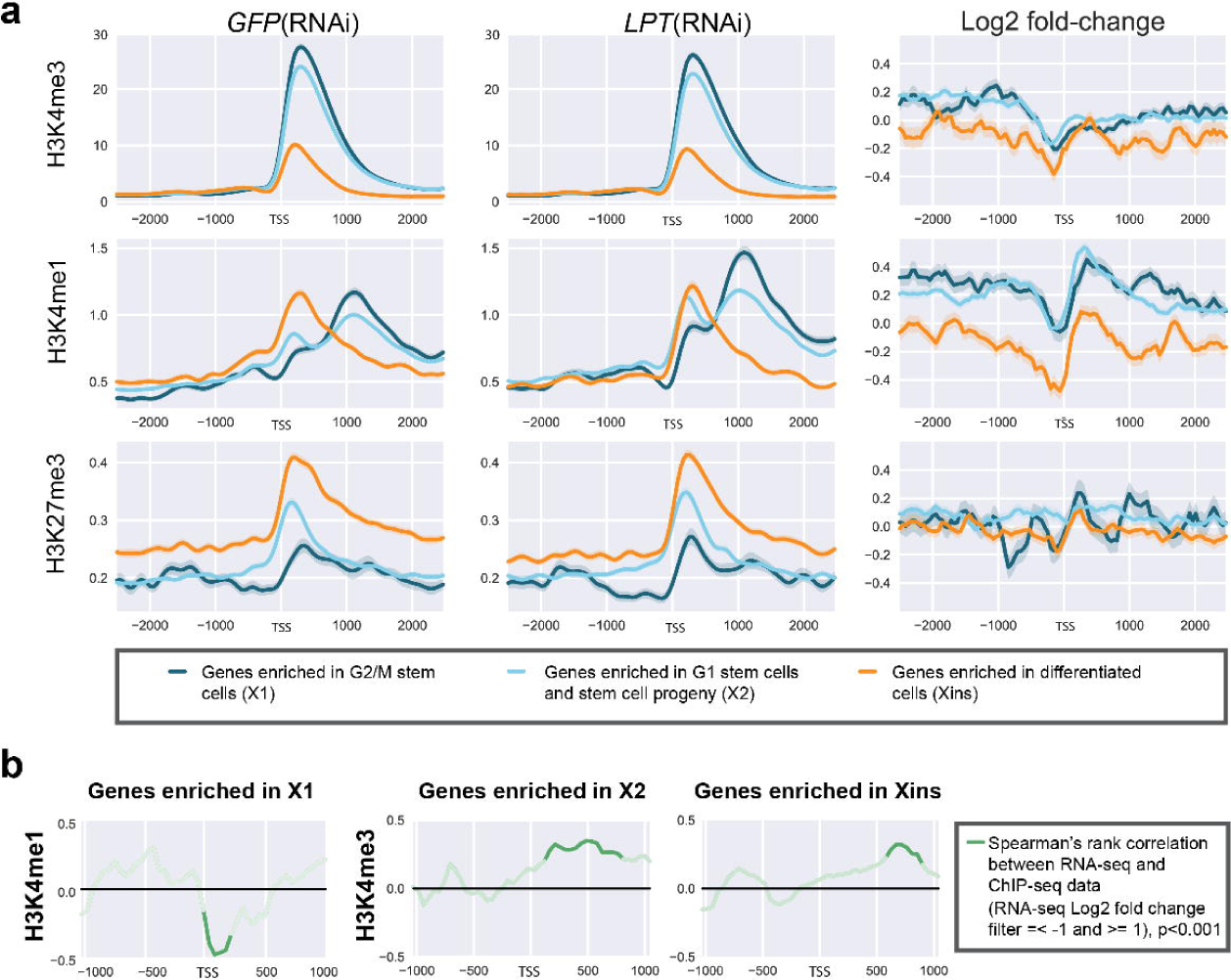
*LPT*(RNAi) is mainly manifested in changes in H3K4me1 and H3K4me3 around the TSS in G2/M (X1) stem cells. **(a)** Graphs presenting the average read coverage across the genome for H3K4me3, H3K4me1 and H3K27me3 after performing ChIP-seq on X1 (G2/M stem cells). The graphs are centered on the TSS (showing 2 kb upstream and downstream) and the data is normalised to *Drosophila* S2 signal spike-in. The input coverage is subtracted. Log2 fold change graphs are also shown for each histone modification, where signal above zero shows increase following *LPT*(RNAi) and signal below zero represents a decrease. Three colours are used for different gene classes - dark blue (genes enriched in G2/M stem cells, i.e. X1), light blue (genes enriched in G1 stem cells and stem cell progeny, i.e. X2), orange (genes enriched in differentiated cells, i.e. X ins). Standard deviation is shown by a faded colour around each line. **(b)** Spearman’s rank correlation between changes in RNA-seq signal following *LPT*(RNAi) and H3K4me1 or H3K4me3 ChIP-seq signal for the region around the TSS of genes from different enrichment classes (only examples where a significant correlation exists are shown). The green line shows a correlation where RNA-seq fold change data was filtered for Log2 fold changes =<-1 and >= +1. Faded areas of the lines represent results not significant at p<0.001, while darker colours represent results significant at p<0.001.

We next looked specifically at the promoter histone methylation status of those genes whose transcript levels were affected by *LPT*(RNAi). For genes with enriched expression in NBs, we observed a significant inverse correlation between expression following *LPT*(RNAi) and the amount of TSS-proximal H3K4me1 (**Figure 8b**). This indicates that *LPT*(RNAi) leads to a reduction of this repressive mark at some loci and an up-regulation of the cognate transcript expression in stem cells, consistent with the role of H3K4me1 as a repressive mark. For mis-regulated genes not normally enriched in X1 NBs, we observed instead a positive correlation between changes in transcriptional expression (upregulation) and changes in H3K4me3 levels at the TSS and gene bodies (**Figure 8b**).

Overall, our data suggest that reductions in H3K4me1 following *LPT*(RNAi) cause up-regulation of some of the stem cell genes implicated by RNA-seq data from *LPT*(RNAi) animals. Our data are consistent with MLL3/4’s known role in H3K4 methylation and identify gene expression profiles following *LPT*(RNAi) that are broadly correlated with the amount of H3K4me1 and H3K4me3 at gene promoters.

### *LPT*(RNAi) leads to up-regulation of known and putative oncogenes and down-regulation of tumour suppressors

After observing the global changes in expression and histone modification patterns following *LPT*(RNAi), we wanted to investigate individually mis-regulated genes that could be major contributors to the differentiation and outgrowth phenotypes (**Figure 9 and 10**). Within our list of up- and down-regulated genes, we saw mis-regulation of tumour suppressors, oncogenes and developmental genes (**Supplementary Table 1**). Detailed inspection of changes in promoter patterns of H3K4 methylation revealed example loci where changes in methylation status were both consistent and inconsistent with changes in transcript levels (**Figure 9a,10a, Supplementary Figure 7a, 9a**). For example, we find that the up-regulated expression of the planarian orthologs of the transcription factors *Elf5* and *pituitary homeobox (pitx)* is associated with increased levels of TSS-proximal H3K4me3 signal following *LPT*(RNAi) (**Figure 9a**). Furthermore, up-regulation of some X1-enriched genes, such as *pim-2-like*, is associated with a decrease in H3K4me1 signal on the TSS, consistent with alleviated repression (**Figure 9a**). On the other hand, some transcriptional changes following *LPT*(RNAi), such as the down-regulation of *Ras-responsive element-binding protein 1 (RREBP1)*, are not correlated with the expected alterations in histone modification patterns on promoters (**Figure 10a**). Such examples could potentially represent secondary (not related to histone modifications) or enhancer-dependent changes in the *LPT*(RNAi) phenotype. In the absence of similar RNA-seq/ChIP-seq data in mammals, our data provide an important insight beyond the deep evolutionary conservation of MLL3/4 function.

**Figure 9.**
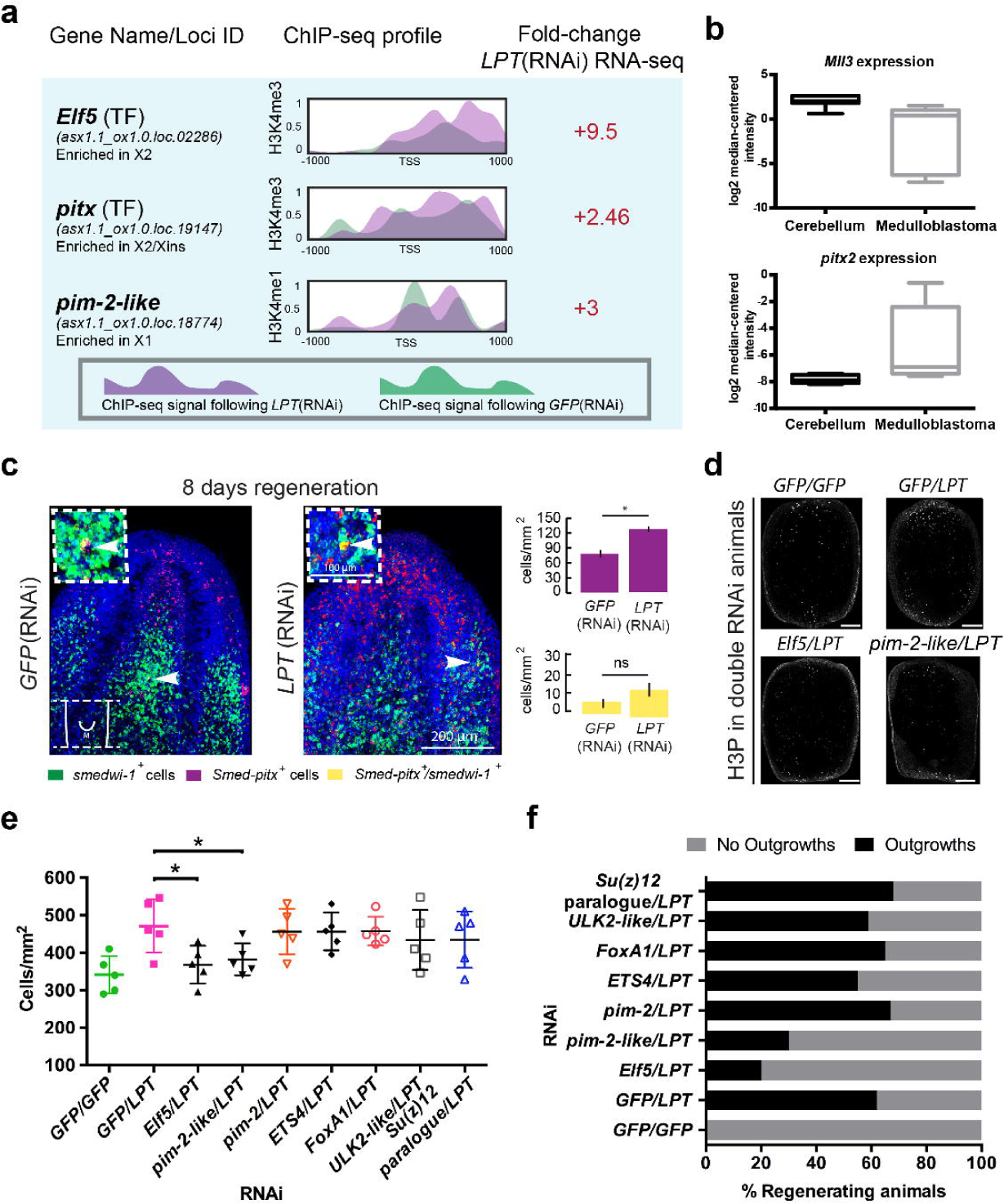
Double knockdown with *Elf5* or *pim-2-like* alleviates the *LPT*(RNAi) over-proliferation and outgrowth phenotype. **(a)** Examples of genes significantly (p<0.05) up-regulated in G2/M stem cells following *LPT*(RNAi). The ChIP-seq profile for H3K4me3 or H3K4me1 in the 2 kb region around the TSS of each gene is presented. Purple colour represents normalised signal following *LPT*(RNAi) and green colour is used to show the normalised signal following GFP(RNAi). The bold font of the gene names signifies a correlation between the genes’ up-regulation and their respective H3K4me3/me1 profile. ‘TF’ stands for ‘transcription factor’. All three genes show correlation between ChIP-seq profile and up-regulation in RNA-seq data. **(b)** *in silico* analysis (www.oncomine.org; t-test, p<0.0001) of Mll3 and pitx2 expression in normal tissue (cerebellum) and cancer tissue (medulloblastoma). **(c)** *pitx* and *smedwi-1 in situ* hybridization at 8 days of regeneration of middle pieces following *LPT*(RNAi). White arrows show double-positive cells. 2-tailed t-test used for analysis, n=10, *p<0.05. **(d)** Representative examples of mitotic cells (labeled by anti-H3P antibody) in double RNAi condition at 48h post amputation. **(e)** Graph showing number of mitotic cells in double RNAi animals at 48h post amputation. Each dot represents average number of mitotic cells in single worm (n=5). Lines and error bars indicate mean and SD. Student’s t test: *p<0.05. **(f)** Graph showing percentage quantification of double knockdown regenerates developing outgrowths.

**Figure 10.**
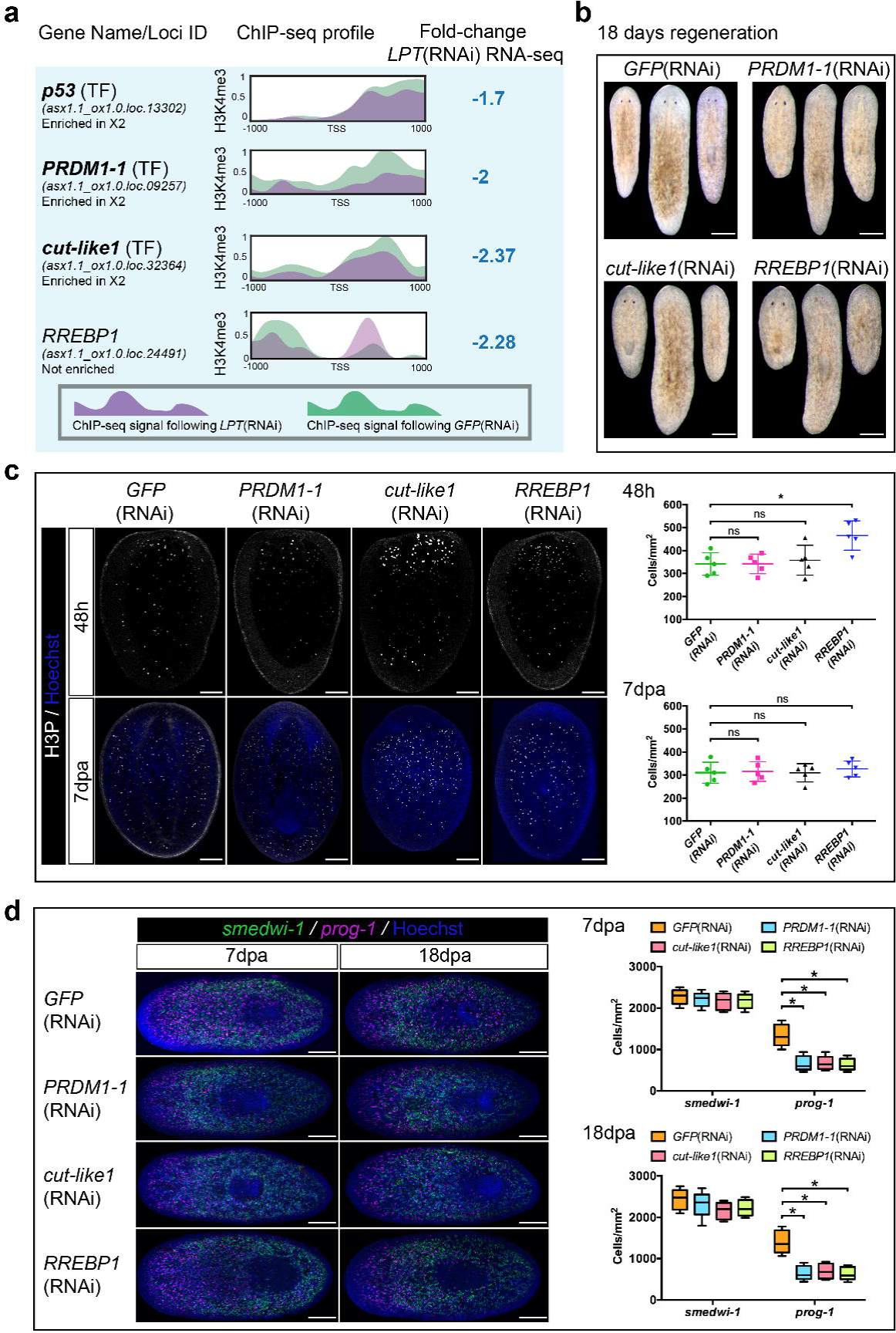
*LPT*(RNAi) down-regulates the expression of genes involved in stem cell proliferation and differentiation. **(a)** Examples of genes significantly (p<0.05) down-regulated in G2/M stem cells following *LPT*(RNAi). The ChIP-seq profile for H3K4me3 in the 2 kb region around the TSS of each gene is presented. Bold font of a gene name illustrates an example where there is a correlation between H3K4me3 profile and down-regulation in RNA-seq data. ‘TF’ stands for ‘transcription factor’. **(b)** Representative bright field images of 18 day regenerating animals following different RNAi. *PRDM1-1* (RNAi), *cut-like1* (RNAi) and *RREBP1* (RNAi) animals show defective anterior regeneration compared to *GFP*(RNAi) animals. Scale bar: 200 µm. **(c)** Representative images of mitotic cells (labeled by anti-H3P antibody) in different RNAi animals at 48h and 7 days post-amputation. Scale bar: 100 µm. Graphs show mitotic cell quantification. Each dot represents average number of mitotic cells in single worm (n=5). Lines and error bars indicate mean and SD. Student’s t test: *p<0.05. **(d)** Representative images showing stem cells *(smedwi-1*^*+*^) and early epidermal progeny *(prog-1+*) at 7 and 18 day regenerating animals from different RNAi conditions. Graphs show quantification of stem cells and early epidermal progeny (n=5). Lines and error bars indicate mean and SD. Student’s t test: *p<0.05.

While mis-regulation of well-known oncogenes and tumour suppressors, like *p53* (**Figure 10a**), would be expected to broadly correlate with mis-regulation of *Mll3/4* in cancer gene expression datasets, shared correlative expression changes for some selected genes not previously associated with *Mll3/4* loss of function provide some independent evidence of a conserved regulatory program. In the Pomeroy Brain Oncomine dataset^59^, *Mll3* and *p53* expression levels were both significantly down-regulated (**Supplementary Figure 10**), supporting the functional link between the two genes established by previous work^32^ and emerging in our study. We identified more *LPT*(RNAi) mis-regulated genes, which have some known association with growth and/or cancer, but have not been previously implicated in *Mll3/4* loss of function phenotypes. A surveillance of the Brune^60^ and Compagno^61^ lymphoma Oncomine datasets demonstrates that these genes are mis-regulated in different cancers in a similar manner to the mis-regulation we observe in planarians stem cells as a consequence of a decreased *LPT* expression (**Supplementary Figure 10**).

One gene of interest in this data set was the planarian *pitx* gene ortholog. In planarians, *pitx* is expressed in the serotonergic neuronal precursor cells^62,63^ and is required for their differentiation. Thus, *pitx* is not directly implicated in planarian stem cell proliferation, but rather in differentiation. Nonetheless, *pitx’s* up-regulation was of great interest to us since in human medulloblastomas down-regulation of *Mll3* and over-expression of *pitx2* are co-occurrences (Pomeroy Brain Oncomine dataset^59^ ((www.oncomine.org)) (**Figure 9b**). To investigate the cellular basis for *pitx* overexpression, we performed FISH for this gene in *LPT*(RNAi) animals. We observed an accumulation of pitx-positive cells in *LPT*(RNAi) regenerates (**Figure 9c**). Given that production of terminally differentiated serotonergic neurons is decreased (**Figure 3d**), the increase of pitx-positive cells following *LPT*(RNAi) marks the accumulation of serotonergic neuronal precursors that fail to differentiate. We conclude that planarian LPT normally regulates pitx-mediated differentiation of serotonergic neurons and that regulation of *pitx* is a possible example of a conserved feature of MLL3/4 function that is mis-regulated in some cancer types^59^.

### Double RNAi experiments allow proof of principle functional validation of overexpressed LPT target genes

In the planarian model, up-regulated genes in our data set provided the opportunity to identify genes whose overexpression contributes to the *LPT/Mll3/4* loss of function phenotype using double RNAi experiments. In addition to the transcription factor *E74-like factor 5 (Elf5)*, we observed up-regulation of planarian orthologs of other developmental or cancer associated genes. Among them were orthologs of the serine/threonine kinase oncogene *pim-2* (two genes called *Smed-pim-2* and *Smed-pim-2-like)*, a paralog of the epigenetic regulator *Suppressor of zeste (Su(z)12)*, the transcription factors *ETS4* and *FoxA1* and an ULK2-like serine/threonine protein kinase (**Figure 9a, Supplementary Figure 7a**). Our ChIP-seq data suggested *pim-2, ETS4 and ULK2-like* are either indirectly regulated or regulated by LPT at enhancers, while *pim-2-like, FoxA1, Su(z)12 paralog* and *Elf5* appear to be a direct target of LPT as their increased expression was associated with either reduced H3K4me1 or increased H3K4me3 signal at promoters (**Figure 9a, Supplementary Figure 7a**).

We decided to focus on these genes because they all have a known role in a wide range of cancers. Overexpression of the cell fate decisions determinant *Elf5* is a known driving force behind breast cancer progression and metastasis^64^. The PIM family of proteins is involved in the integration of growth and survival signals^65^ and their overexpression has been associated with hematological malignancies and solid tumours^66^. Inhibition of PI M2 is a promising avenue for the treatment of multiple myeloma^65^. Su(z)12 is part of the ubiquitous polycomb repressive complex 2 (PRC2) and is overexpressed in numerous cancers, including gastric cancer where Su(z)12 levels were associated with increased metastasis and unfavorable survival prognosis^67^. The ETS family of transcription factors has demonstrated significant involvement in all stages of tumorigenesis^68^, while FoxA1 is known as ‘pioneer transcription factor in organogenesis and cancer progression’^69^. Finally, ULK2 is an autophagy regulator overexpressed in prostate cancer cells^70^. This selected panel of genes represents some of the best candidates for major effects amongst those genes with significant up-regulation in expression following *LPT*(RNAi).

In order to test whether the up-regulated expression of any of these genes is a potentially significant contributor to the *LPT*(RNAi) outgrowth phenotype, we performed *LPT*(RNAi) rescue experiments in the form of double RNAi knockdowns (**Figure 9d-f, Supplementary Figure 7**). At 48 hours postamputation, *LPT*(RNAi) regenerates have a significantly increased NB proliferation (Figure 4a, b) and so do GFP/*LPT*(RNAi) double knockdown animals (Figure 9d,e). Whereas *pim-2/*LPT*(RNAi), ETS4/*LPT*(RNAi), FoxA1/*LPT*(RNAi), ULK2-like/LPT(RNAi*) and *Su(z)12 paralogue/LPT(RN*Ai) regenerates still have elevated NB proliferation, both *pim-2-like/*LPT*(RNAi)* and Elf5/*LPT*(RNAi) regenerates have a significantly decreased NB proliferation compared to GFP/*LPT*(RNAi) (**Figure 9d,e and Supplementary Figure 7b**). Furthermore, not only did *pim-2-like/LPT*(RNAi) and Elf5/*LPT*(RNAi) regenerating animals show improved blastema formation (**Supplementary Figure 7c**), but also less than half as many animals in these two conditions went on to form outgrowths compared to GFP/*LPT*(RNAi) (**Figure 9f**), demonstrating a rescue of the outgrowth phenotype. Importantly, individual knockdown of *Elf5* and *pim-2-like* did not lead to regenerative, proliferation or outgrowth-related defects (**Supplementary Figure 8**). These findings suggest that the up-regulation of both *pim-2-like* and *Elf5* is specifically involved in driving the *LPT*(RNAi) outgrowth phenotype and demonstrate the utility of our data for validating the role of MLL3/4 targets.

### *LPT*(RNAi)’s differentiation defects can be explained by down-regulated transcription factors

We chose a selection of transcription factors down regulated by **LPT*(RNAi)* for further investigation to see if they might contribute to the observed **LPT*(RNAi)* phenotype. *LPT*(RNAi) leads to the down-regulation of the tumour suppressor *p53, PRDM1-1* and *cut-like1* in X1 stem cells (**Figure 10a**). These genes’ down-regulation is associated with a decrease in H3K4me3 levels on their promoters. On the other hand, RREBP1’s down-regulation, does not correlate with the amount of H3K4me3 on its promoter. While the function of *p53* in planarians has already been described^71^, the role of *PRDM1-1, cut-like1* and *RREBP1* remain unexplored in planarian biology. Knockdown of each of these three genes resulted in impaired regeneration, characterized mainly by the inability to form eyes (**Figure 10b**). RREBP1(RNAi) resulted in significantly increased proliferation compared to control, while *cut-like1* (RNAi) and *PRDM1-1* (RNAi) animals did not show a significant change in mitotic cell numbers (**Figure 10c**). All three knockdown conditions showed a decreased number of *prog-1* -positive epidermal progenitors, but intact *smedwi-1* -positive cell numbers (**Figure 10d**). These findings suggest that the observed differentiation phenotypes are not associated with an inability to maintain the stem cell pool following knockdown. Instead, stem cells seem to be restricted in their ability to differentiate correctly.

Our data suggest that decreased expression of *PRDM1-1, cut-like1* and *RREBP1* in *LPT*(RNAi) animals could be contributing to the differentiation defects seen after perturbation of LPT/Trr/MLL3/4 function, and that downregulation RREBP1 may be contributing to the early over proliferation of stem cells observed in **LPT*(RNAi).*

## Discussion

In mammals, *Mll3* and *Mll4* have been implicated in different malignancy landscapes^24^, with clear evidence for tumour suppressor roles in mammalian systems^26,32,72^. However, relatively little is known about how these effects are mediated. Our study demonstrates that loss of function of the planarian *LPT* (an *Mll3/4* ortholog) also results in the emergence of an outgrowth phenotype characterized by differentiation and proliferation defects. Our work also shows that LPT, TRR-1 and TRR-2 control differentiation to form the gut, eyes, brain, and pharyngeal cellular lineages, Future work in planarians combining ChIP-seq and RNA-seq will allow closer investigation of these and other epigenetic effects on stem cell differentiation.

We found that clusters of mitotic cells preceded the appearance of outgrowths in *LPT*(RNAi) regenerating animals, possibly pre-empting where the outgrowths would subsequently form. The observation of clusters of cells and the formation of outgrowths in some, but not all, RNAi animals is evidence of heterogeneity in stem cell responses to *LPT*(RNAi). This may reflect the stochastic nature of the broad genome wide epigenetic changes mediated by MLL3/4 proteins that will lead to variability between cells after knockdown, such that only some NBs cycle out of control and cause outgrowths after the initial proliferative peaks associated with regeneration. A contributory cause to outgrowth formation in addition to proliferation could be failure of NBs to differentiate appropriately and instead continue to cycle at inappropriate positions. We also observed that outgrowth tissue contained different classes of stem cells. Among these stem cells, the presence of *sigma* NBs, thought to include truly pluripotent stem cells^55^, is of particular significance. When mis-regulated, these cells could share fundamental similarities with cancer stem cells (CSCs), thought to be founder cells in human malignancies^73^. CSCs have been described as one of the main factors in cancer aggressiveness and resistance to treatment^74^. Studying such cells in a simple *in vivo* stem cell model, provided by the planarian system, should bring further insight into important control mechanisms that are mis-regulated in different cancers. Our work here provides a useful example of this approach.

Our data suggest that LPT regulates expression of genes across cell types, including some genes with enriched expression in stem cells. Genes with significant expression differences following *LPT*(RNAi) were mostly associated with cell proliferation, differentiation and metabolic processes. A subset of mis-regulated genes where RNA-seq and ChIP-seq data correlate is likely a direct consequence of *LPT*(RNAi) affecting promoter histone methylation status. Genes with altered expression where there is no such correlation, may represent indirect (secondary) changes or, alternatively, may have enhancers that have altered histone modifications as a result of *LPT*(RNAi). Future work will develop the use of planarians as a model of epigenetic gene regulation and it should also be possible to study enhancer function and evolution.

Amongst mis-regulated genes, we saw many tumour suppressors with reduced expression and oncogenes with increased expression. We also found a number of genes, including the transcription factor *pitx*, which were similarly mis-regulated in *LPT*(RNAi) planarians and human cancers with reduced *Mll3* expression. Together, these data suggest that, as well as physiological function in controlling stem cell proliferation, there may be deep regulatory conservation of MLL3/4 function in animal stem cells. One advantage of our approach is that we were able to sample expression and histone states in NBs at an early time point before tumours formed. This could be an advantage for the identifying targets that act early to drive hyperplasia, rather than later secondary regulatory changes.

As proof of principle that genes mis-regulated by *LPT*(RNAi) directly contributed to the phenotype, we performed double RNAi experiments. Planarian homologs of the oncogene *pim-2*, called *Smed-pim-2* and *Smed-pim-2-like*, that were overexpressed in stem cells following *LPT*(RNAi), were chosen as likely candidates, based on previous data on the roles of these genes from mammals^66-70^. We found that double RNAi with *pim-2-like*, was able to ameliorate *LPT* loss of function over-proliferation and outgrowth phenotypes induced by *LPT*(RNAi). In addition, double knockdown with the breast cancer oncogene *Elftf*^*64*^ resulted in an even more dramatic rescue of the LPT loss of function phenotype. This provides strong support for the hypothesis that the over-expression of these two genes was significant in driving stem cell hyperplasia. Future work can now study how these genes function in stem cells and why overexpression leads to overproliferation.

We also identified downstream candidates that could be contributing to the lack of differentiation phenotype following *LPT*(RNAi). Knockdown of *PRDM1-1, cut-like* and *RREBP-1* (down-regulated in *LPT*(RNAi) animals) indicated their mis-regulation might contribute to the decreased epidermal differentiation observed following *LPT* knockdown.

Togtether these experiments, demonstrate the value of our approach to identify potential downstream targets and implicate novel regulatory interactions driving the *Mll3/4* loss of function phenotype. These targets can now be tested for conservation in mammalian experimental systems.

Overall, our work shows how perturbation of a conserved physiological role of LPT leads to mis-regulation of genes well-known to control cell proliferation, causing hyperplasia and tumours in planarians. We find other genes that are mis-regulated in planarians that are also similarly mis-regulated in cancer expression studies that have reduced *Mll3* expression. Some of these, like *pitx*, may represent deeply conserved regulatory interactions. In the absence of similar RNA-seq/ChIP-seq data in mammals, our data provide an important insight into *Mll3/4* loss of function, as well as revealing a deep evolutionary conservation in animal stem cells. These findings demonstrate the strength of the planarian system for understanding fundamental animal stem cell biology and the potential for investigation of epigenetic mechanisms in stem cells.

## Methods

### Animal husbandry

Asexual freshwater planarians of the species S. *mediterranea* were used. The culture was maintained in 1x Montjuic salts water^75^. Planarians were fed organic calf liver once a week. After every feeding, the water was changed. Planarians were starved for 7 days prior to each experiment. They were also starved throughout the duration of each experiment.

### RNAi

Double-stranded RNA (dsRNA) was synthesized from DNA fragments cloned in pCRII (Invitrogen) or pGEM-T Easy (Promega) vectors. T7 (Roche) and SP6 (NEB) RNA polymerases were used for transcription of each strand. The two transcription reactions were combined upon ethanol precipitation. RNA was denatured at 68 °C and re-annealed at 37 °C. Quantification was performed on a 1% agarose gel and Nanodrop spectrophotometer.

For single RNAi experiments a working concentration of 2 µg/µl was used. For double RNAi, each gene’s RNA was at a concentration 4 µg/µl, resulting in solution concentration of 2 µg/µl.

DsRNA was delivered via microinjection using Nanoject II apparatus (Drummond Scientific) with 3.5’’ Drummond Scientific (Harvard Apparatus) glass capillaries pulled into fine needles on a Flaming/Brown Micropipette Puller (Patterson Scientific). Each animal received around 100 nl dsRNA each day. *H2B*(RNAi) was performed for three consecutive days, as per Solana et al.’s (2012) protocol^7^. For single and double *LPT, trr-1* and *trr-2* knockdown, a course of 7 days of microinjections was performed (3 consecutive days + 2 days rest + 4 consecutive days). Set1(RNAi) and utx(RNAi) were performed for 4 consecutive days. For all other single and double knockdowns, a course of 10 days of microinjections was performed (3 consecutive days + 4 days rest + 3 consecutive days).

Primers used for amplification of DNA for dsRNA synthesis can be found in **Supplementary Table 2**.

### In situ hybridization

RNA probes labeled with digoxigenin and fluorescein were generated via antisense transcription of DNA cloned in PCRII (Invitrogen) or PGemTEasy (Promega) vector. *In situ* hybridization was performed as described in King and Newmark’s protocol^76^ for most fluorescent experiments. For *LPT, trr-1, trr-2, sigma, zeta* and *gamma* fluorescent *in situ* procedures, a pooled probes method was used, as described in van Wolfswinkel et al.^55^. Colorimetric *in situ* hybridization procedures were performed as described in Gonzalez-Estevez et al.^77^. Primers used for amplification of DNA for RNA probe synthesis can be found in (**Supplementary Table 2**).

### Immunohistochemistry

Immunohistochemistry was performed as described in Cebria and Newmark^78^. Antibodies used were: anti-H3P (phosphorylated serine 10 on histone H3; Millipore; 09-797; 1:1000 dilution), anti-VC1 (kindly provided by Prof. Hidefumi Orii (check title); 1:10000 dilution), anti-SMEDWI-1 (kindly provided by Prof. Jochen Rink; 1:500 dilution), anti-SYNORF-1 (3C11; Developmental Studies Hybridoma Bank; 1:50 dilution), anti-acetylated tubulin (Developmental Studies Hybridoma Bank; 1:200 dilution).

### Imaging and image analysis

Colorimetric images were taken on Zeiss Discovery V8 (Carl Zeiss) microscope with a Canon EOS 600D or Canon EOS 1200D camera. Fluorescent images were taken on either Inverted Olympus FV1000 or FV1200 Confocal microscope. Cells were counted via Adobe Photoshop CS6 or FIJI software and the count was normalized to image area in mm^2^.

### Flow cytometry

A modified version of Romero et al.’s^79^ planarian FACS protocol was used, as described in Kao et al.^9^. A FACS Aria III machine equipped with a violet laser was used for the sort. BD FACSDiva and FlowJo software was used for analysis and gate-setting.

### Western blot

2xLaemmli buffer (Sigma Aldrich), 1M DTT and cOmplete protease inhibitors (Roche) were used for protein extraction from 10-15 animals per condition. Protein extract was quantified with Qubit Protein Assay kit (Thermo Fisher Scientific). NuPAGE Novex 4-12% Bis-Tris protein gels (Thermo Fisher Scienitific) were used, followed by a wet transfer in a Mini Trans-Blot Electrophoretic Transfer Cell machine. Ponceau S (Sigma Aldrich) whole-protein stain was used prior to antibody incubation. The antibodies used were: anti-H3 (unmodified histone H3; rabbit polyclonal; Abcam; ab1791; 1:10000 dilution), anti-H3K4me3 (rabbit polyclonal; Abcam; ab8580; 1:1000 dilution), anti-H3K4me1 (rabbit polyclonal; Abcam; ab8895; 1:1000 dilution), anti-H3K27me3 (mouse monoclonal; Abcam; ab6002; 1:1000 dilution), anti-mouse IgG HRP-linked antibody (Cell Signalling; 7076P2), anti-rabbit IgG HRP-linked antibody (Cell Signalling; 7074P2). Western blot experiments were done to validate the specificity of the histone modification antibodies used for ChIP-seq (**Supplementary Figure 11**). The rationale behind these experiments was to knock down a methylase *(set1)*, part of a methylase complex *(LPT)* or a demethylase *(utx)* known to affect H3K4 and H3K27 methylation levels and to observe whether global H3K4me1, H3K4me3 and H3K27me3 levels would change in the expected way.

### ChIP-seq

600,000-700,000 planarian x1 cells were FACS-sorted (using 3-day knockdown regenerates) in PBS and pelleted at 4 °C. During the pelleting, S2 cells were added (corresponding to roughly 15% of the number of planarian x1 cells) for the purpose of downstream data normalisation^58^. Samples were then processed as described in Kao et al. (2017)^9^. The process is summarized in **Supplementary Figure 12**. The libraries were sequenced on an Illumina NextSeq machine. Three biological replicates were prepared. The raw reads are available in the Short Read Archive (PRJNA338116).

### RNA-seq

300,000 x1 NBs were FACS-sorted in RNALater (Ambion) from knockdown animals at 3 days of regeneration. Cells were pelleted at 4 °C and Trizol-based total RNA extraction was performed. The amount of total RNA used for each library preparation was 0.8-1 µg. Illumina TruSeq Stranded mRNA LT kit was used for library preparation. The kit instructions were followed. Libraries were quantified with Qubit, Agilent Bioanalyzer and KAPA Library Quantification qPCR kit. Samples were sequenced on an Illumina NextSeq machine. Two biological replicates were prepared. The raw reads are available in the Short Read Archive (PRJNA338115).

### ChIP-seq data analysis

ChIP-seq reads were trimmed with Trimmomatic 0.32^80^ and aligned to the S. *mediterranea* SmedGD asexual genome 1.1^81^ and *D. melanogaster* genome r6.10^82^ with BWA mem 0.7.12. Picard tools 1.115 was used to remove read duplicates after mapping. Python scripts were used to filter and separate out read pairs belonging to either genome. ChIP-seq coverage tracks were then generated and normalized according to *Orlando et al*. in order to account for any technical variation between samples^58^. For more in-depth methods, including code, refer to **Supplementary Note 1**.

### RNA-seq data analysis

Raw reads were trimmed with Trimmomatic 0.32^80^ and pseudo-aligned to a set of asexual genome annotations described in Kao et al. (2017) with Kallisto 0.42^83^. Differential expression was subsequently performed with Sleuth 0.28.1^84^. For more in-depth methods, including code, refer to **Supplementary Note 1**.

### Statistical methods

Wherever cell number was compared between experimental condition and control, a 2-tailed t-test assuming unequal variance was used. Each legend states the number of specimens per condition, where relevant. Bar graphs show the mean average and the error bars are always Standard Error of the Mean.

For analysis of RNA-seq data, Wald’s test (as part of the Sleuth^85^ software) was used for assessing differential expression. Spearman’s rank correlation was used for assessing the correlation between RNA-seq and ChIP-seq data. Hypergeometric tests were used for assessing gene enrichment in the RNA-seq data.

### Data availability

The ChIP-seq and RNA-seq datasets are deposited in the Short Read Archive with accession numbers: PRJNA338116 and PRJNA338115 respectively).

The ‘Pomeroy Brain’ dataset^59^ from the oncomine database (https://www.oncomine.com) was used for assessing expression level of pitx2 and *Mll3* in human medulloblastoma versus normal cerebellum. All other data availability is either within the article (and its supplementary information).

## Authors’ contributions

AAA, PA and YM conceived and designed the study. YM and PA performed the experiments. DK performed the bioinformatics analyses. SH participated in the optimization of the ChIP-seq protocol. AGL provided technical support. FJH performed initial work on the project, including generating the first *LPT*(RNAi) results. NK helped with sigma, zeta and gamma *in situ* hybridization experiments. YM, PA and AAA wrote the manuscript.

## Acknowledgements

We thank past and present members of the AA lab for comments on the manuscript.

## Supplementary Information

**Supplementary Figure 1. Structure and function of COMPASS and COMPASS-like core proteins.**

**(a)** Schematics of the core subunits of the COMPASS and the two COMPASS-like complexcorresponding to different protein domaines in mammals are presented with coloured boxes corresponding to different protein domains – RRM1 (RNA-recognition motif), N-SET, SET, CXXC (zinc finger), PHD (Plant Homeodomain fingers), zf (PHD-like zinc finger), FYRN (Phenylalanine/Tyrosine rich N-terminus domain), FYRC (Phenylalanine/Tyrosine rich C-terminus domain), purple stars signifying nuclear receptor recognition motifs. Dashed vertical line represents proteolytic cleavage. **(b)** As in **(a)**, but in fruitfly. **(c)** Proposed mechanisms of action of each core complex subunit. COMPASS complex – 1) performing H3K4 trimethylation on TSS of most actively transcribed genes and 2) depositing H3K4me2 on the gene bodies of actively transcribed genes. MLL1/2/Trithorax COMPASS-like complex – 1) a role in transcriptional activation of Hox genes via trimethylating H3K4 on TSS of their promoters and 2) MLL2 is involved in trimethylation of H3K4 on TSS of bivalent promoters. MLL3/4/LPT/Trr – 1) role in hormone-dependent transcription – when the Nuclear Receptor protein (NR) is bound to the DNA Hormone Response Element (HRE) upon Hormone Ligand (HL) detection, MLL3/4/LPT/Trr complex binds the nuclear receptor and serves as its coactivator via trimethylating H3K4 and promoting active transcription on selected loci; 2) a switch between inactive and active enhancer states where MLL3/4/LPT/Trr complex deposits H3K4me1 on both active and inactive enhancers; upon UTX recruitment, it demethylates H3K27me3 and allows for CBP/p300 to acetylate H3K27 and activate the enhancer; 3) a switch between active and inactive promoters – MLL3/4/LPT/Trr complex bound to TSS deposits H3K4me1 on the TSS and around it, leads to repressed transcription of the gene; when H3K4me1 is depleted from the TSS and another complex performs trimethylation of H3K4 on TSS, this is correlated with activated transcription. **(d)** Schematic representation of planarian COMPASS and COMPASS-like core subunits. SMED-LPT (in red) is characterized in the present study. **(e)** Planarian COMPASS and COMPASS-like core subunits’ expression in the three populations of cells sortable by fluorescence-activated cell sorting (FACS) (X1=G2/M stem cells, X2=G1 stem cells and stem cell progeny, X ins=differentiated cells) according to RNA-seq data. **(f)** Known defects after RNAi-mediated knockdown of core COMPASS and COMPASSlike subunits in planarians.

**Supplementary Figure 2. Planarian *MII3/4* genes are expressed in neoblasts and neoblast progeny and colocalise with each other.**

**(a)** Protein alignment of conserved regions of COMPASS-like families’ core proteins. Asterisks indicate complete conservation in all sequences, while black boxes are drawn around areas of conservation specific to the MLL3/4/Trithorax-related family. Colours represent similarity of amino acids. The image was produced using MEGA.5.2 software.

**(b)** Bright field images of head, middle and tail pieces following *trr-1*(RNAi), *trr-2*(RNAi) or control *GFP*(RNAi) at day 8 of regeneration. Yellow arrows point towards the regenerative defects – smaller blastema, delayed eye formation or posterior bloating.

**(c)** Head, middle and tail pieces following *trr-1*(RNAi), *trr-2*(RNAi) or control *GFP*(RNAi) at day 14 of regeneration.

**(d)** Central nervous system (CNS) maintenance and recovery at 8 days of middle piece regeneration, as labeled by CNS-specific anti-SYNORF-1 antibody, following *trr-1*(RNAi) or *trr-2*(RNAi).

**(e)** Bright field images of head, middle and tail pieces at 3 days of regeneration following *GFP/GFP*(RNAi), *trr-1/trr-2*(RNAi), *GFP/trr-2*(RNAi), *GFP/trr-1*(RNAi) and *GFP/LPT*(RNAi). Red arrows point towards outgrowths.

**(f)** Survival curve of middle regenerating pieces in different RNAi conditions. The *GFP/GFP*(RNAi) line overlaps with *GFP/trr-1*(RNAi) and *GFP/trr-2*(RNAi). n=10.

**Supplementary Figure 3. *Trr-2*(RNAi) regenerating animals produce less GABAergic and dopaminergic neurons.**

**(a)** Quantification of the number of GABAergic neurons (labeled by *GAD*), **(b)** dopaminergic neurons (labeled by *TH*), **(c)** serotonergic neurons (labeled by *TPH*), **(d)** acetylcholinergic neurons (labeled by *chat*) and **(e)** early (labeled by *prog-1*) and late (labeled by *AGAT-1*) epidermal stem cell progeny at 8 days of regeneration of tail or middle pieces following trr-1(RNAi) or trr-2(RNAi). 2tailed t-test used for analysis, n=10, *p<0.05. Error bars represent Standard Error of the Mean (SEM).

**Supplementary Figure 4. *Sigma, zeta* and *gamma* neoblast numbers are unchanged following *LPT*(RNAi).**

**(a)** FISH showing cells in 8 days of regenerating animals following *LPT*(RNAi) labeled by the *sigma pool* of RNA probes (*Soxp1, Soxp2*) and *smedwi-1.* White arrows point towards *sigma* neoblasts (double-positive for *sigma pool* and *smedwi-1).*

**(b)** FISH showing cells in 8 days of regenerating animals following *LPT*(RNAi) labeled by the *zeta pool* of RNA probes (*zfp-1, Soxp3, egr-1*) and *smedwi-1.* White arrows point towards *zeta* neoblasts (double-positive for *zeta pool* and *smedwi-1).*

**(c)** FISH showing cells in 8 days of regenerating animals following *LPT*(RNAi) labeled by the *gamma pool* of RNA probes (*gata4/5/6, hnf4*) and *smedwi-1.* White arrows point towards *gamma* neoblasts (double-positive for *gamma pool* and *smedwi-1).*

**(d-g)** Graphs showing quantification of sigma **(d)**, zeta **(e)**, gamma **(f)** and total *smedwi-1*^*+*^ **(g)** neoblasts in 8-day regenerating animals following *LPT*(RNAi). Each dot represents average number of cells in a single worm (n=5). Lines and error bars indicate mean and SD. Student’s t test: *p<0.05.

**Supplementary Figure 5. *LPT*(RNAi) results in disorganized outgrowth-focused expression of epidermal precursor markers, epithelial disarray, hypertrophy and changes of nuclear morphology.**

**(a)** Anterior part (containing an outgrowth) of a tail piece at 18 days of regeneration following *LPT*(RNAi) labeled with *prog-1* and *AGAT-1* epidermal precursor markers. ‘CG’ stands for ‘cephalic ganglia’.

**(b)** The epidermal layer (stained with Hoechst 33342) of a tail piece at 10 days of regeneration following *LPT*(RNAi) compared to control.

**(c)** Graph showing increase in nuclear area following *LPT*(RNAi). 2-tailed t-test used for analysis, n=20, ***p<0.001.

**(D)** Image showing changes in nuclear morphology of epidermal cells in 10-day regenerating animals following *LPT*(RNAi). Nuclei were stained with Hoechst 33342. Yellow arrows point towards misshapen nuclei.

**Supplementary Table 1. Differentially expressed loci following *LPT*(RNAi).**

Each row represents one locus that was differentially expressed with a p-value less than 0.05 and fold change <-1.5 or >1.5. The Wald’s test (as part of the Sleuth software) was used for assessing differential expression. The top BLAST hit (with e-value) and the common model organism top BLAST hit is also provided for each locus.

**Supplementary Figure 6. Histone modification ChIP-seq profiles at promoter-proximal regions of different classes of genes.**

**(a-c)** Images show histone modification patterns for H3K4me3 **(a)**, H3K4me1 **(b)** and H3K27me3 **(c)** respectively. ChIP-seq signal is shown in black. Three classes of genes are presented – enriched >50% in X1 (G2/M stem cells) shown by dark blue, enriched >50% in X2 (G1 stem cells and stem cell progeny) shown in light blue, enriched >50% in X ins (differentiated cells) shown in orange. Histone modification graphs are centered on the Transcriptional Start Site (TSS) with 2.5 kb shown upstream and downstream.

**Supplementary Figure 7. Simultaneous knockdown of LPT with Elf5 or pim-2 like results in partial recovery of the *LPT*(RNAi) regenerative phenotype.**

**(a)** Examples of genes significantly (p<0.05) up-regulated in G2/M stem cells following *LPT*(RNAi). The ChIP-seq profile for H3K4me3 in the 2 kb region around the TSS of each gene is presented. Purple colour represents normalised signal following *LPT*(RNAi) and green colour is used to show the normalised signal following GFP(RNAi). ‘TF’ stands for ‘transcription factor’. Bold font of a gene name illustrates an example where there is a correlation between H3K4me3 profile and up-regulation in RNA-seq data.

**(b)** Representative examples of mitotic cells (labeled by anti-H3P antibody) in double RNAi condition at 48h post amputation.

**(c)** Bright field images showing partial recovery in 10 day regenerating animals following *Elf5/LPT(*RNAi) and *pim-2 like/LPT(R*NAi) compared to *GFP/LPT*(RNAi). Other genes screened failed to recover the regeneration defects.

**Supplementary Figure 8. RNAi of genes up-regulated following *LPT*(RNAi) did not result in defects in regeneration or stem cell proliferation.**

**(a)** Bright field images showing regeneration following RNAi of different genes up-regulated in *LPT*(RNAi).

**(b)** Images showing mitotic cells (labeled by anti-H3P antibody) in regenerating animals following knockdown of different genes up-regulated in *LPT*(RNAi). Graph showing mitotic cell quantification following gene knockdowns.

**Supplementary Figure 9. Knockdown of some genes down-regulated following *LPT*(RNAi) does not result in regeneration defects.**

**(a)** Examples of genes significantly (p<0.05) down-regulated in G2/M stem cells following *LPT*(RNAi). The ChIP-seq profile for H3K4me3 and H3K4me1 in the 2 kb region around the TSS of each gene is presented. Purple colour represents normalised signal following *LPT*(RNAi) and green colour is used to show the normalised signal following GFP(RNAi). ‘TF’ stands for ‘transcription factor’. Bold font of a gene name illustrates an example where there is a correlation between H3K4me3 profile and down-regulation in RNA-seq data.

**(b)** Bright field images of 18-day regenerating animals following RNAi of different genes down-regulated in *LPT*(RNAi). Scale bar: 200 µm.

**(c)** Representative examples of mitotic cells (labeled by anti-H3P antibody) at 48h and 7 day post amputation in regenerating animals following knockdown of different genes down-regulated in *LPT*(RNAi). Scale bar: 100 µm. Graphs show the quantification of mitotic cells. Each dot represents average number of mitotic cells in a single worm (n=5). Lines and error bars indicate mean and SD. Student’s t test was used for analysis.

**(d)** Representative FISH images showing stem cells (*smedwi-1*^*+*^) and early epidermal progeny (*prog-1^+^*) at 7 day regenerating animals in different RNAi conditions. Scale bar: 100 µm. Graph shows the quantification of *smedwi-1*^*+*^ and *prog-1^+^* cells (n=5). Lines and error bars indicate mean and SD. Student’s t test used for analysis.

**Supplementary Figure 10. Mis-regulation of genes following *LPT*(RNAi) correlates with mis-regulation in human cancers where *MII3* levels are decreased.**

*In silico* analysis (www.oncomine.org; t-test, p<0.0001) of Mll3, IGFALS, p53, ULK2, TBX1 and cut-like expression in normal tissue (cerebellum or different B-lymphocyte types) and cancer tissue (medulloblastoma, follicular lymphoma or Hodgkin’s lymphoma). Positive and negative numbers next to gene names indicate up- or down-regulation in *LPT*(RNAi) X1 RNA-seq respectively.

**Supplementary Table 2. Primer sequences**. All primers are given in 5’->3’ orientation. ‘F’ and ‘R’ stand for ‘forward’ and ‘reverse’ primer respectively.

**Supplementary Figure 11. The histone modifications antibodies used for ChIP-seq experiments are specific.**

**(a)** Western blot with anti-H3K4me1 and loading control anti-H3 (unmodified histone H3) on protein lysate from *GFP*(RNAi) and *LPT*(RNAi) animals.

**(b)** Western blot with anti-H3K4me3 and loading control anti-H3 (unmodified histone H3) on protein lysate from *GFP*(RNAi) and set1(RNAi) animals.

**(c)** Western blot with anti-H3K27me3 and loading control anti-H3 (unmodified histone H3) on protein lysate from *GFP*(RNAi) and *utx(RNAi*) animals.

**Supplementary Figure 12. Summary of planarian ChIP-seq procedure.**

Three day-regenerating planarians were dissociated into single cells. Cells were stained with Hoechst 34580 and Calcein AM in order to visualize cell populations according to nuclear size and cytoplasmic complexity. The X1 (G2/M) stem cells (magenta) were sorted and mixed with 4% *Drosophila* S2 cells. Cells were crosslinked with 1% Formaldehyde and sonicated. Immunoprecipitation with anti-H3K4me3, anti-H3K4me1 and anti-H3K27me3 antibodies followed. Samples were reverse-crosslinked and libraries were prepared using NEBNext Ultra II library preparation kit.

**Supplementary Note 1. Supplementary Python Notebook.**

Provides details on the ChIP-seq and RNA-seq bioinformatics analyses.

